# CAI-Vax: Computational Design of Personalized mRNA Cancer Vaccine Constructs After Antigen Selection

**DOI:** 10.64898/2026.10.20.760576

**Authors:** Jalen Cai, Yulin Chen, Yi Sun, IpYan He, Lareina Yuan, Eileen Liu, Wilson Ho-Man Lee, Zhipeng Lu

**Author notes:** Correspondence: Jalen Cai (; alternate).

## Abstract

CAI-Vax (Cancer Antigen Individualized Vaccine Design) arranges selected antigens and generates synonymous coding sequences for personalized mRNA vaccine constructs. We evaluated 357 vaccine-candidate peptides from 31 patients across four published cohorts. At the primary cumulative MHCflurry percentile-rank threshold of ≤2%, HLA-aware search reduced predicted junction-spanning peptide-allele binder occurrences by 3.49 per junction relative to amino-acid-pattern optimization (paired Hodges–Lehmann estimate; nominal 95% interval 2.75–4.48; Holm-adjusted P = 7.45 x 10^-9^). All 31 primary paired differences and the estimated effects in all 16 threshold-by-normalization settings favored HLA-aware search within the MHCflurry model used for optimization.

Among 25 patients with an exact solution for the additive search objective, single-start greedy search found the minimum in 5 patients and the best individual heuristic in 16. Selecting each patient’s best result across all tested heuristics increased this to 19. MHCflurry and NetMHCpan EL identified identical tied best-plan sets in 6 of 31 patients after rescoring six frozen plans per patient. In a retrospective internal analysis, repeat-aware synonymous encoding reduced maximum 10-mer multiplicity in every patient, with mean codon-use-preference losses of 0.0041–0.0055 across cohorts; however, it performed worse than matched random encoding for 6-mers. These findings demonstrate that HLA-guided construct optimization can reduce predicted junctional binding within the optimization model, although the preferred antigen arrangement depends on the prediction model used. Experimental studies are needed to determine the biological and manufacturing relevance of these computational differences.

## Introduction

Cancer is not a uniform disease, even among patients with the same histologic diagnosis. Individual tumors differ in their somatic mutations, transcriptional states, clonal architecture, and mechanisms of immune evasion. The clinical success of immune-checkpoint blockade established that endogenous T-cell responses can produce durable tumor control in some patients, but the benefit remains incomplete and heterogeneous across tumor types and individuals.^1^ Somatic mutations can create altered protein sequences that are absent from normal tissues. When these proteins are expressed, processed into peptides, and presented by human leukocyte antigen (HLA) molecules, the resulting neoantigens may be recognized by T cells. This tumor specificity makes neoantigens attractive targets for personalized cancer vaccines.^2^ Early clinical studies showed that individualized neoantigen vaccination can induce mutation-specific T-cell responses, supporting the feasibility of patient-specific vaccine development.^3,4^

Messenger RNA is well-suited to individualized vaccines because antigen sequences can be changed for each patient while the broader production platform remains constant. mRNA can encode multiple antigens, is translated in the cytoplasm, and produces transient expression without requiring nuclear entry. Advances in nucleoside modification, purification, sequence engineering, and delivery have improved translation and reduced undesirable innate immune sensing.^5–7^ Personalized RNA and mRNA vaccine studies have induced mutation-specific T-cell responses in melanoma, pancreatic cancer, and other solid tumors.^4,8,9^ Randomized phase 2b testing in resected melanoma has further supported the clinical development of individualized mRNA neoantigen therapy in combination with immune-checkpoint blockade.^10^

Tumor and matched-normal DNA sequencing identify somatic variants, tumor RNA sequencing measures mutant-transcript expression, and HLA typing defines the patient’s alleles. Candidate peptides can then be assessed with predictors such as NetMHCpan and NetMHCIIpan, and subsequently prioritized with workflows such as pVACtools.^11,12^ Experimental confirmation and expert review remain important during neoantigen candidate selection.^13–15^

After the antigen set has been chosen, the antigens must be ordered and, when applicable, separated by linkers to form a vaccine construct. Different arrangements generate different peptides spanning the junctions between adjacent antigens. Some of these peptides may be predicted to bind the patient’s HLA alleles, creating junctional epitopes that were not part of the selected antigen set. pVACvector addresses this problem by searching antigen order and spacer configurations.^12^ Here, we evaluate antigen arrangement and linker selection while treating each patient’s antigen set as a fixed upstream input (Fig. 1).

**Figure 1.**
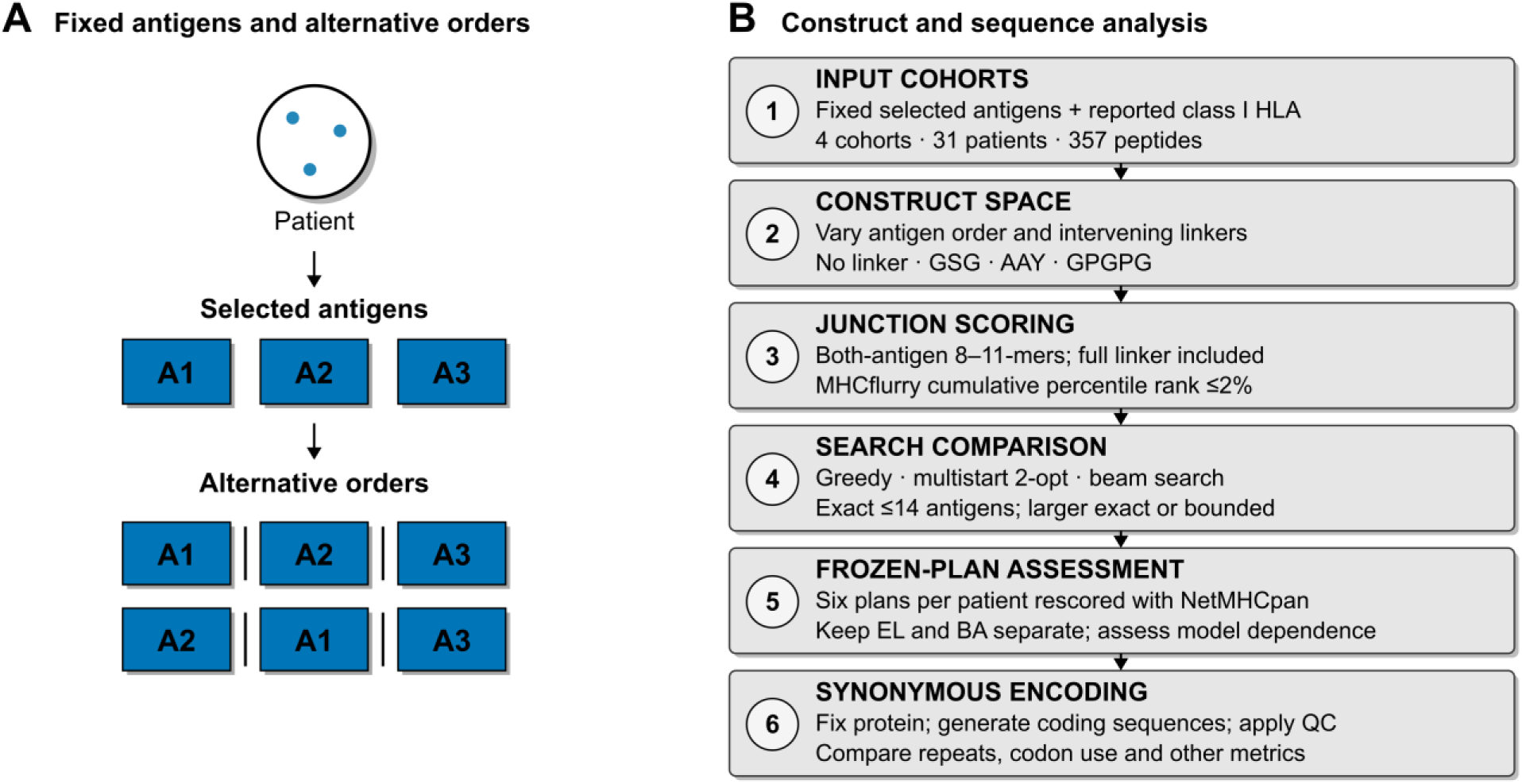
Construct design after antigen selection. (A) Each patient contributes a fixed antigen set. Reordering changes antigen junctions while preserving antigen identities. Evaluated peptides were 8–11 amino acids long and included at least one amino acid from each adjacent antigen and the entire linker, when present. (B) CAI-Vax evaluates these peptides, compares heuristic search with exact or bounded optimization, freezes plans for cross-predictor comparison, and generates synonymous coding sequences subject to quality-control checks.

Encoding the resulting protein cassette introduces a second design problem. Because the genetic code is degenerate, many synonymous coding sequences can produce the same amino-acid sequence while differing in codon usage, GC composition, CpG and UpA content, homopolymers, and predicted RNA folding. These properties can interact, such that improving one sequence metric may worsen another.^16–18^ A useful workflow must therefore distinguish structural requirements such as preserving the intended amino-acid sequence and reading frame from comparative objectives whose biological importance remains uncertain.

We developed CAI-Vax to address these post-selection design decisions. The workflow searches antigen orders and linker assignments using amino-acid features and predicted HLA class I binding. It then generates synonymous coding sequences, applies structural quality-control checks, and compares sequence metrics with matched random encodings and fixed-rule baselines.

CAI-Vax evaluation was conducted using fixed antigen sets from four published patient cohorts, including a multisector glioblastoma cohort.^19^ We asked whether HLA-aware search reduced predicted junctional binding, how closely heuristic searches approached the minimum of the specific objective, whether construct rankings transferred between HLA predictors, and what trade-offs arose during synonymous encoding. All reported outcomes were computational design metrics; antigen selection itself was outside the scope of the study.

## Methods

### Study design and scope

We retrospectively evaluated CAI-Vax using fixed antigen sets from published patient cohorts. Each patient contributed one independent observation to patient-level statistical comparisons. Outcomes included construct-design metrics, coding-sequence metrics, and search performance.

### Software and reproducibility

CAI-Vax was implemented in Python. The accompanying analysis archive preserves the source snapshot, input checksums, configurations, random seeds, and retained result tables. Supplementary Methods describe the analysis version and reproduction boundaries; the archive provides commands and dependency records.

A separately documented predictor reproduction matched the retained frozen-plan counts for the historical inputs. After correcting the PGV001 patient mapping, we newly searched and scored the two affected patients and retained the 29 unaffected records. Supplementary Methods distinguish the historical reproduction from the current 186-plan analysis and report predictor versions, model checks and remaining provenance limits.

### Input sources and cohort definitions

CAI-Vax accepts pVACseq tables, peptide FASTA files, pVACvector-style FASTA files, and antigen manifests.^12^ An HCC1395 cell-line example was used to test input handling and was excluded from patient analyses. We analyzed four published cohorts: multisector glioblastoma (4 evaluable patients), hepatocellular carcinoma (9 patients), melanoma NeoVax cohort (6 patients), and PGV001 (12 evaluable patients).^3,19–21^ Glioblastoma was selected as the development cohort; the remaining cohorts were locked before retrospective evaluation. The analysis included 31 total patients. Patients were deemed evaluable if at least one non-empty antigen sequence containing supported standard amino-acid symbols was available. Of 13 PGV001 patient identifiers in the source tables, PID-002 was excluded because its available supplementary entries did not contain extractable peptide sequences. In Table S3, the label PGV-019 was normalized to PID-019 before joining to Table S2 HLA metadata. The previous parser failed to recognize this prefix and assigned ten PID-019 peptides to PID-017. Correcting the mapping separates the two patients with ten peptides each and leaves the total of 357 evaluable peptides unchanged. Published immune-response, manufacturing, and clinical annotations were retained.

### Input harmonization

Source records were converted to a common patient-antigen table containing the sequence, patient and antigen identifiers, source location, and available HLA and upstream-selection metadata. Invalid or duplicate records and failed joins were handled using recorded rules. Each construct search used antigens from one patient only.

### Cassette assembly and amino-acid junction features

For each patient, the selected antigens were preserved as intact amino-acid sequences and assembled into candidate translated cassettes. A construct consisted of an ordered list of antigens and one linker assignment at each adjacent seam. The available linker options were GSG, AAY, GPGPG, and no linker. Uniform-linker searches used the same option at every junction, whereas mixed-linker searches allowed the option to vary among junctions. Every construct was checked to confirm preservation of all source antigen sequences and the expected number of seam assignments.

Each junction was evaluated with separate amino-acid features, including homopolymer runs, basic- and hydrophobic-residue runs, low complexity, repeats, and cassette length. In the corrected comparisons, the amino-acid-pattern objective minimized the edge-additive number of flagged junctions. Cost ties among linker options were resolved first by shorter linker length and then by the declared order: no linker, GSG, AAY, GPGPG. Amino-acid-pattern and HLA-aware searches were compared under matched search configurations. The Supplementary Methods define the feature thresholds, treatment of intentional linker repeats, linker tie-breaking rule, and search configurations.

### Patient-specific HLA class I junction assessment

Patient class I alleles were taken from the source records without imputation and checked for predictor support. At each junction, we enumerated peptides 8–11 amino acids long that contained at least one residue from each flanking antigen, including the complete intervening linker when present. MHCflurry 2.2.1 affinity percentile ranks were obtained for unique peptide-allele pairs within each junction.^22^ Missing or unsupported predictions were reported separately. Within-antigen peptides, linker-only peptides, and peptides containing residues from only one flanking antigen were outside this endpoint.

The strong-binder threshold was an inclusive percentile rank of ≤0.5%, and the primary cumulative threshold was ≤2%. The cumulative ≤2% count included predictions meeting the ≤0.5% threshold. Numeric percentile ranks were compared inclusively with each threshold. MHCflurry burden was the sum of qualifying unique peptide-allele occurrences across junctions; consequently, the same peptide-allele pair could contribute at two different junctions. We reported raw counts, counts per junction, counts per supported allele, and counts per 100 construct-wide unique peptide-allele opportunities. The per-100 measure is an occurrence-to-unique-opportunity ratio and need not be a percentage. Sensitivity analyses crossed thresholds of 0.5%, 1%, 2%, and 5% with these four reporting scales.

### Frozen independent-predictor rescoring

We rescored six frozen construct plans for each patient with NetMHCpan, producing 186 patient-plan combinations.^23^ The current table combines 174 unchanged records from 29 patients with 12 newly scored records for corrected PGV001 patients PID-017 and PID-019. The new scoring used the documented licensed 4.2e installation. Antigen order, linker assignments, HLA alleles, peptide lengths and thresholds were fixed before rescoring. Eluted-ligand and binding-affinity ranks were analyzed separately.

NetMHCpan counts used a construct-wide union of eligible peptide-allele pairs. Two frozen uniform-linker plans repeated peptide sequences across junctions; both had zero recorded MHCflurry counts at ≤0.5% and ≤2%. The other 184 plans had no cross-junction duplicates. Union deduplication therefore leaves the frozen comparisons unchanged at these two thresholds. Statistical analyses retained each patient’s six correlated plans within the same cluster.

### Cross-predictor agreement and reproducibility

We reconstructed the same six plans for each patient and repeated MHCflurry scoring. Reproduction required identical integer binder counts and matching construct fingerprints. Cross-predictor analyses compared tied best-plan sets, rank reversals, regret, and non-dominated candidates separately from the optimization objective.

MHCflurry selected the minimum raw cumulative ≤2% counts across the six candidates, with exact equality defining ties. NetMHCpan EL evaluated each candidate using its cumulative count per 100 candidate-specific peptide-allele opportunities. Pessimistic regret was the largest EL score among the MHCflurry-tied minima minus the lowest EL score among all six candidates. Because selection and evaluation used different scales and candidate-specific denominators, this regret measure reflects both predictor disagreement and differences in eligible opportunities.

### Construct search and search-correctness analysis

For each directed antigen pair, scalar search chose the linker with the minimum edge cost. The HLA edge cost was the raw cumulative ≤2% MHCflurry binder-occurrence count. Ties were resolved first by shorter linker length, then by the declared order: no linker, GSG, AAY, GPGPG. The resulting path objective was additive across junctions.

Held–Karp dynamic programming found the exact minimum of this additive Hamiltonian-path objective for patients with no more than 14 antigens.^24^ Larger instances were analyzed using a mixed-integer linear program with subtour-elimination constraints, implemented with SciPy and HiGHS.^25,26^ Heuristics included single-start and all-start greedy search, 32 random restarts followed by 2-opt, and beam search with widths 16, 32, 64, and 128.

Adaptive search used batches of multistart 2-opt and stopped after reaching its combinatorial lower bound, completing two consecutive batches without strict improvement, or reaching the runtime check. Exact or bounded solutions were used only afterward to assess the adaptive result. The Supplementary Methods specify the batch schedule, seed, solver limits, evaluation counts, and measured path-search runtime.

### Multi-objective Pareto search

A custom label-setting implementation retained non-dominated alternatives for two minimized edge-additive objectives: amino-acid flags and MHCflurry binder occurrences. A plan dominated another if it was no worse in both objectives and strictly better in at least one. Uncapped searches were exact within the configured antigen limit; capped searches were approximations. Crowding-distance truncation used the NSGA-II rule to preserve objective extremes.^27^ The recurrence, label limits, and validation cases are given in Supplementary Methods.

### Synonymous coding-sequence generation

After a protein cassette was fixed, synonymous coding sequences were generated in the DNA alphabet and interpreted as mRNA after T-to-U conversion. Available strategies included random synonymous encoding, high-frequency-codon encoding, GC-balanced encoding, low-CpG encoding and low-UpA encoding, homopolymer-avoidance encoding, and the balanced CAI-Vax encoder. The balanced encoder selected synonymous codons using explicit terms for deviation from the target GC fraction of 0.52, newly created CpG and UpA dinucleotides, homopolymer burden, and human codon-use preference. Every strategy was required to reproduce the intended amino-acid sequence.

When a supplied cassette did not begin with methionine, the encoder added an initiating methionine and recorded the resulting sequence as the intended translated product.

Structural validation compared each coding sequence with that recorded product and verified the terminal-stop convention. Initiating methionine was added in 29 of the 31 source-order GSG benchmark cassettes; the two exceptions were HCC patients N09 and N31.

### Hard constraints and sequence-feasibility metrics

Structural failures always invalidated a candidate, including empty coding sequences, unsupported nucleotide characters, length not divisible by three, premature internal stop codons, missing or incorrect terminal-stop structure, translation mismatch, unintended amino-acid additions or deletions, and coding-sequence length above the configured maximum of 12,000 nucleotides.

Candidate sequences that passed structural checks were characterized by whole-sequence GC content, minimum and maximum sliding-window GC content, mean codon usage, CpG and UpA fractions, maximum nucleotide homopolymer run, repeated k-mer multiplicity, motif burden, and optional RNA-folding features. Default feasibility thresholds included a whole-sequence GC interval of 30-75%, a 50-nucleotide sliding window with 20-85% GC limits, a maximum homopolymer threshold of 10 nucleotides, and repeated 10-mer multiplicity tracking. These thresholds were warnings by default and could be promoted to hard failures through configuration. Minimum-free-energy and local folding features were calculated with the ViennaRNA Package.^28^ Sequence-feasibility and manufacturability-proxy metrics were treated as engineering heuristics.

### Repeat-aware synonymous optimization

The repeat-aware encoder began with the standard CAI-Vax coding sequence and performed constrained synonymous local descent. Accepted substitutions preserved the intended protein, maintained whole-sequence GC content within 30-75%, did not increase the starting homopolymer length or exceed ten nucleotides, and limited the reduction in mean codon-use preference to 0.02.

Objective weights were chosen separately for each held-out cohort using the other three cohorts. The 4,000-evaluation work threshold was calibrated using the historical 30-patient inputs and held fixed for the corrected 31-patient rerun. Thus, only objective-weight selection was isolated by cohort; the analysis was a retrospective internal evaluation. The Supplementary Methods provide the complete objective, selection rule, evaluation counts, and matched random-encoding baselines.

### Matched internal baselines

Matched random encodings sampled synonymous codons for the same intended protein. The original composite-score benchmark generated 500 random encodings per patient; the repeat-length baseline analysis used 120. Fixed-rule baselines included high-frequency, GC-balanced, and low-repeat encoders. Generator rules, favorable directions, tie treatment, invalid-draw handling, and seeds are detailed in the Supplementary Methods.

### Statistical analysis

Patients were the independent statistical units. Paired differences were summarized with the Hodges–Lehmann estimator, calculated as the median of the Walsh averages.^29^ The primary nominal 95% interval used Walsh order statistics selected by a normal approximation to the signed-rank distribution and was not bootstrapped. For samples smaller than six, the implementation returned the observed range of Walsh averages, not 95% confidence limits.

Two-sided Wilcoxon signed-rank tests used the exact conditional distribution of the positive rank sum, obtained by enumerating sign assignments through dynamic programming.^30^ True zero differences were removed, and equal absolute nonzero differences received average ranks. Normalized patient differences were calculated from integer counts and exact rational denominators, avoiding ties determined by display rounding or floating-point subtraction. Two-sided P values were twice the smaller inclusive tail probability, capped at one; all-zero comparisons received P=1. The test assumes independent patient differences and sign symmetry under the null hypothesis. Holm adjustment was applied within the declared contrast families.^31^ This post-review recalculation retained the original contrast definitions and multiplicity groups. Patient-cluster bootstrap intervals resampled patients while retaining correlated plans together; codon-use-loss intervals bootstrapped patient means.^32^ Patient proportions used Clopper–Pearson intervals.^33^ One-sided random-encoding and sign tests were kept separate from two-sided patient tests. The Supplementary Methods specify the interval procedures, resampling counts, seeds, and comparison families.

### Analysis reporting

Runs recorded inputs and exclusions, structural checks, component metrics, prediction completeness, baseline comparisons, search status, and configuration. Missing data, failed analyses, and negative findings were retained in the reports.

## Results

### Cohort and construct overview

We analyzed 357 vaccine-candidate peptides from 31 patients across four published cohorts (Table 1). Glioblastoma was the development cohort; the other cohorts were evaluated retrospectively after their inputs were locked. Construct strategies were compared under matched search configurations. All synonymous coding sequences preserved the intended amino-acid sequence and reading frame. GC content, homopolymers, repeated sequences, and codon use were evaluated as separate metrics.

**Table 1.** Retrospective cohort composition and analysis role.*

| **Cohort** | **Evaluable patients** | **Selected peptides** | **Analysis role** |
| --- | --- | --- | --- |
| Glioblastoma | 4 | 36 | Development cohort |
| Hepatocellular carcinoma | 9 | 107 | Locked retrospective |
| Melanoma NeoVax | 6 | 98 | Locked retrospective |
| PGV001 | 12 | 116 | Locked retrospective |
| **Total** | **31** | **357** |  |

\* GBM, glioblastoma; HCC, hepatocellular carcinoma. PGV001 contained 13 source patient identifiers, but PID-002 was excluded because peptide sequences could not be extracted from the available supplementary PDF.

### Corrected HLA-aware construct search

At the primary cumulative ≤2% percentile-rank threshold, HLA-aware mixed-linker search reduced MHCflurry binder occurrences relative to the corrected amino-acid-pattern mixed-linker comparator in all 31 patients. The paired Hodges–Lehmann estimate was 3.49 fewer occurrences per junction (nominal 95% CI 2.75–4.48; Holm-adjusted P = 7.45 x 10^-9^; Fig. 2A). This estimate is the median of the Walsh averages of the paired differences.

**Figure 2.**
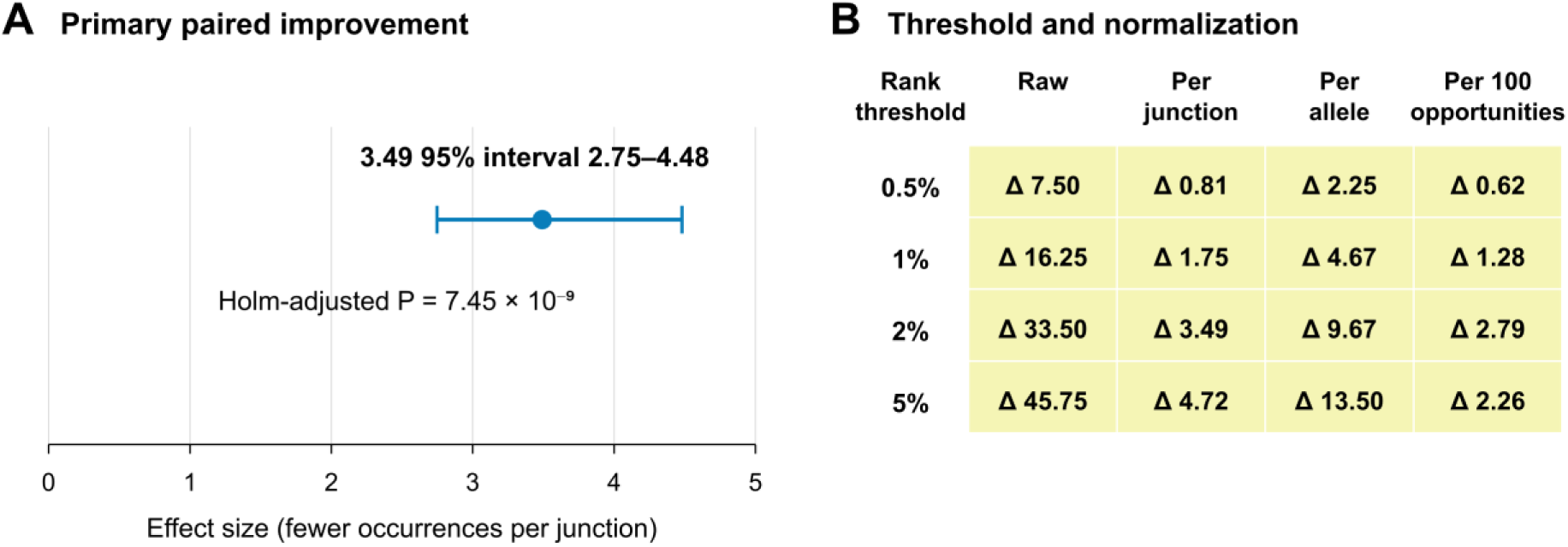
MHCflurry burden under HLA-aware and amino-acid-pattern search. (A) The point shows the paired Hodges–Lehmann estimate for the primary cumulative ≤2% endpoint: 3.49 fewer cumulative binder occurrences per junction with HLA-aware search. The horizontal bar shows the nominal Walsh-based 95% interval of 2.75–4.48. All 31 paired differences favored HLA-aware search. (B) Sensitivity analyses crossed four percentile-rank thresholds with raw, per-junction, per-allele, and per-100-opportunity reporting scales. Each cell shows the Hodges–Lehmann estimate for amino-acid-pattern minus HLA-aware burden. Positive values favor HLA-aware search. All 16 estimates were positive and were Holm-significant within their declared eight-contrast families.

The estimated difference favored HLA-aware search in all 16 combinations of threshold and normalization scale (Fig. 2B; Supplementary Table S1). The statement that all patients improved applies to the primary ≤2% per-junction setting; patient-level directions differed in some sensitivity settings.

### Search correctness and heuristic recovery

Among the 25 patients for whom Held–Karp dynamic programming established the exact minimum of the additive objective, single-start greedy search reached the minimum in 5 and multistart 2-opt, the best individual heuristic, reached it in 16. Selecting the best result obtained for each patient across all tested heuristics increased recovery to 19 of 25. The method giving the best result could differ between patients.

Mixed-integer optimization certified the minimum in 5 of the 6 larger instances. For melanoma patient 2, the best feasible mixed-integer score was 4 and the combinatorial lower bound was 0, leaving an uncertified additive gap of 4. Relative to the predecessor configuration, corrected search reduced the stored score in 21 of 31 patients, with a mean reduction of 4.52 and a maximum reduction of 17 binder occurrences.

### Threshold and normalization robustness

HLA-aware mixed-linker search had a favorable paired estimate in all 16 threshold-by-normalization settings when compared with the corrected amino-acid-pattern mixed-linker plan, source-order GSG, and source-order no-linker plans. The uniform HLA-aware versus uniform amino-acid-pattern comparison also favored HLA-aware search in all 16 settings. These contrasts were Holm-significant within their declared families.

Corrected search also had a favorable estimate relative to its predecessor configuration in all settings, reaching Holm significance in 14 of 16 mixed-linker contrasts and 13 of 16 uniform-linker contrasts. Comparisons with source-order AAY and comparisons between HLA-aware mixed plan and the selected optimized uniform plan did not support consistent conclusions across all settings. The Supplementary Results summarize these comparisons; the accompanying source table supplement_hla_all_128_contrasts.tsv provides all 128 contrasts.

### Flexible-linker threshold-dependent negative result

Mixed-linker designs had a lower predicted binder burden than uniform-linker designs at the primary threshold and were significant within the primary test family. The direction reversed at the 5% threshold. The apparent advantage of mixed linkers therefore depended on the selected binding threshold.

### Repeat-aware synonymous encoding

The repeat-aware encoder reduced maximum repeated 10-mer multiplicity in all 31 patients. Mean reductions were 4.0 in glioblastoma, 5.3 in hepatocellular carcinoma, 8.3 in melanoma, and 4.9 in PGV001 (Table 2). The glioblastoma comparison was not statistically significant (n = 4; two-sided Wilcoxon P = 0.125). Mean losses in codon-use preference ranged from 0.0041 to 0.0055. Cohort-specific bootstrap intervals excluded zero but remained below the configured loss margin of 0.02 (Supplementary Table S3). Objective weights were selected using the other three cohorts in each leave-one-cohort-out fold. However, the 4,000-evaluation work threshold had been calibrated using the historical 30-patient inputs and held fixed for this corrected rerun. The analysis was therefore a retrospective internal evaluation. Across repeat lengths of 6, 8, 10, 12, 15, and 20 nucleotides, the encoder was optimized for 10-mers; for 6-mers, it performed worse than matched random encoding.

**Table 2.** Repeat-aware encoder versus the starting standard CAI-Vax GSG coding sequence across held-out cohort folds.*

| Cohort fold | n | Mean reduction | HL estimate | Interval | Raw P |
| --- | --- | --- | --- | --- | --- |
| GBM | 4 | 4.0 | 3.75 | 2–7 range | 0.125 |
| HCC | 9 | 5.3 | 5.5 | 3.5–7.5 nominal CI | 0.003906 |
| Melanoma | 6 | 8.3 | 8 | 5.5–11.5 nominal CI | 0.03125 |
| PGV001 | 12 | 4.9 | 5 | 4.5–5.5 nominal CI | 0.0004883 |

\* *Reduction is the decrease in maximum nucleotide 10-mer multiplicity. P values are unadjusted two-sided patient-level signed-rank tests. For the four-patient GBM cohort, the reported interval is the observed Walsh range. Objective weights were selected using the other three cohorts, but the work threshold was calibrated globally. Supplementary Table S3 reports codon-use losses and their patient-mean bootstrap intervals.*

### Pareto analysis

For antigen sets containing no more than 11 antigens, uncapped label-setting search recovered non-dominated alternatives that the earlier approximation had omitted. The exact implementation matched exhaustive enumeration in the small real and synthetic instances tested. Larger searches limited the number of labels per state and therefore returned approximate fronts.

### Adaptive no-oracle search

Adaptive search was run for all 31 patients. A certified offline reference was available for 30; melanoma patient 2 was excluded from this denominator because its reference remained uncertified. Adaptive search matched the certified reference in 16 of the 30 evaluable patients. Eleven of these 16 runs established optimality during deployment by attaining the combinatorial lower bound. Other runs stopped after two consecutive batches without strict improvement or at the runtime check. The schedule permitted no more than 32 random restarts plus one greedy start. Search effort and path-search runtime were recorded separately from predictor-inference time.

### Reproduction of retained outputs

The historical reproduction matched 180 retained MHCflurry burden vectors and 30 original mixed-plan fingerprints for the earlier inputs. The corrected analysis preserves the 29 unaffected patient records and replaces the former pooled PID-017 record with two correctly mapped ten-antigen patients. New inference covered all required peptide– allele pairs for their 12 frozen plans, with no missing predictions or unsupported allele pairs (Fig. 3B). All 31 selected synonymous encodings passed structural checks; the 29 unaffected cases exactly reproduced their earlier sequence hashes, evaluation counts, and repeat reductions.

**Figure 3.**
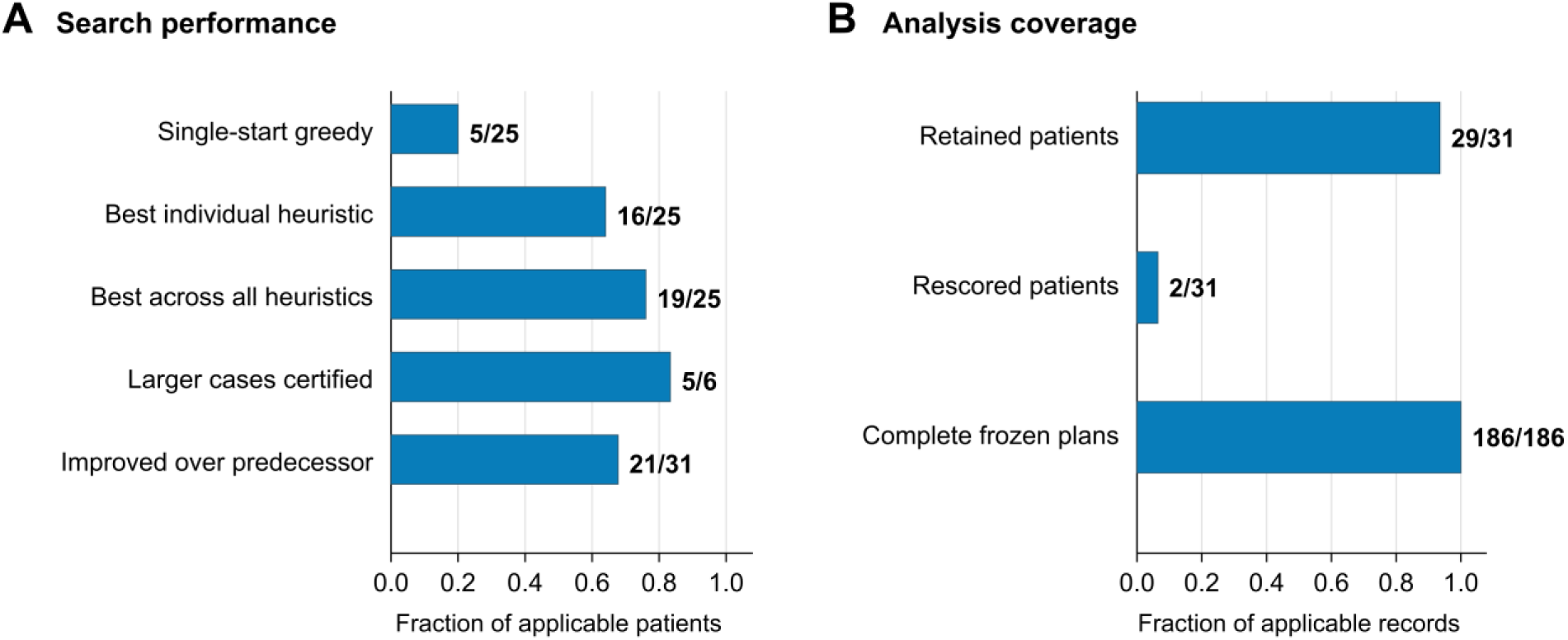
Search performance and analysis coverage. (A) Single-start greedy search reached the exact minimum in 5 of 25 patients and the best individual heuristic in 16. Selecting each patient’s best result across all tested heuristics increased this to 19. Mixed-integer optimization certified 5 of 6 larger cases. Corrected search improved the predecessor score in 21 of 31 patients. (B) Bars show the proportions of patient records retained unchanged (29/31) and newly searched and rescored after the PGV001 mapping correction (2/31), and the proportion of frozen patient-plan records with complete supported peptide-allele predictions (186/186). The shared axis shows proportions; labels give the counts and their denominators.

### Frozen NetMHCpan rescoring and predictor dependence

Across 186 frozen patient-plan observations, the pooled Spearman correlation between MHCflurry and NetMHCpan was 0.871 for EL and 0.719 for BA. These pooled estimates include both between-patient and within-patient differences.

After subtracting each patient’s mean score for each predictor and pooling the 186 residual pairs, the corresponding patient-centered correlations were 0.282 for EL and - 0.191 for BA (Fig. 4A). When correlations were instead calculated separately within each patient, the median was 0.346 for EL and -0.133 for BA among the 30 patients with defined correlations. These three summaries answer different questions; no formal decomposition of between-patient and within-patient variation was performed.^36^

**Figure 4.**
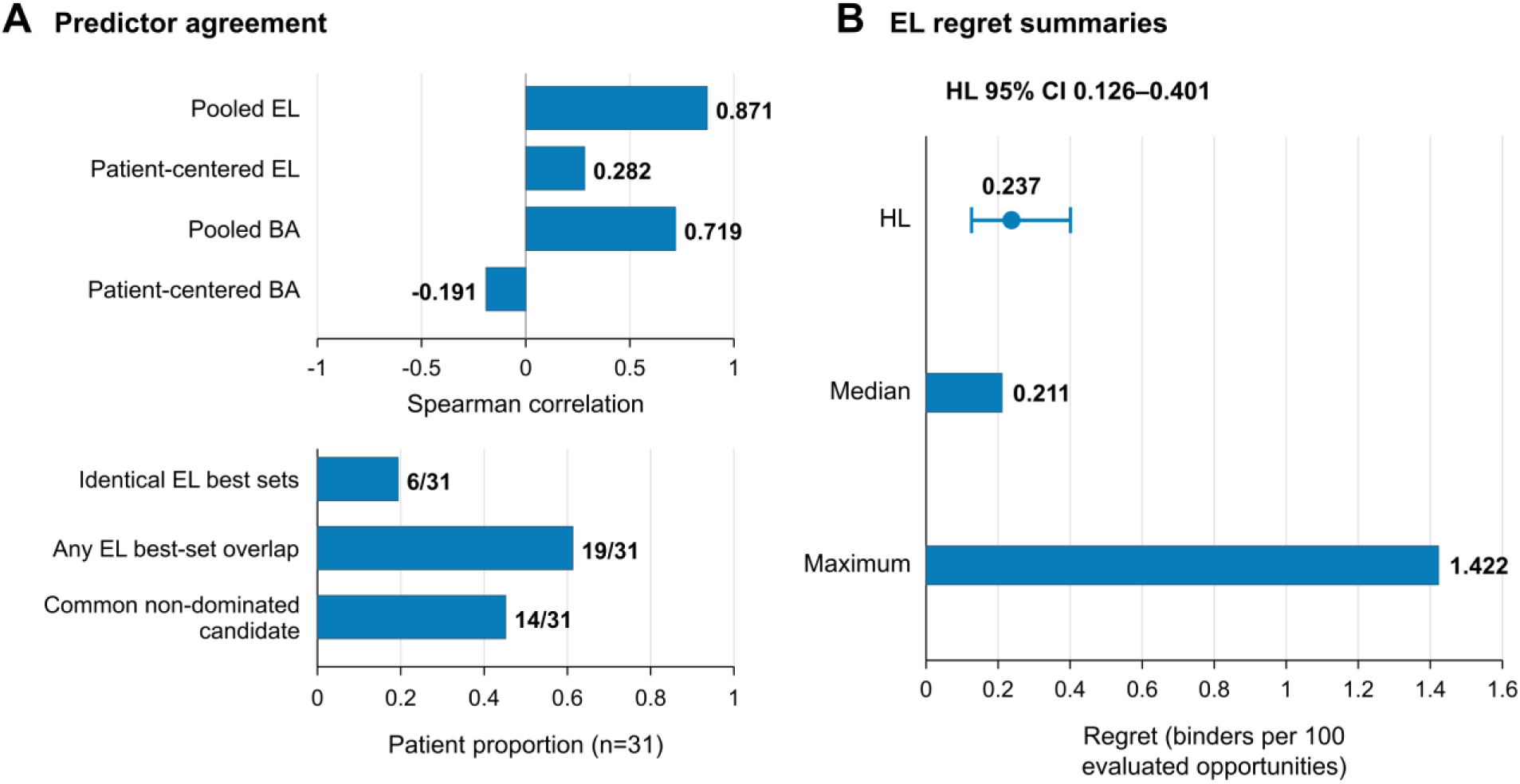
Predictor agreement and EL regret for six frozen plans per patient. (A) Pooled correlations (EL 0.871; BA 0.719) and patient-centered pooled correlations (EL 0.282; BA −0.191) use the correlation axis. The separate proportion axis shows identical EL best sets (6/31), any EL best-set overlap (19/31), and a common non-dominated candidate across three views (14/31). (B) Pessimistic EL regret summaries are HL 0.237 (patient bootstrap 95% interval 0.126–0.401), median 0.211, and maximum 1.422 binders per 100 evaluated opportunities. The HL estimate is shown as a point with its patient-bootstrap interval; horizontal bars show the median and maximum. MHCflurry selection used raw counts.

MHCflurry and NetMHCpan EL identified identical tied best-plan sets in 6 of 31 patients and sets with at least one plan in common in 19 of 31. Under raw-count MHCflurry selection and candidate-normalized EL evaluation, pessimistic regret was zero in 10 of 31 patients. Median regret was 0.211 binders per 100 opportunities; the Hodges– Lehmann estimate was 0.237 (patient bootstrap 95% CI 0.126–0.401), and the maximum was 1.422 (Fig. 4B).

In 14 of 31 patients, at least one frozen candidate was non-dominated in each of three separate two-objective comparisons: amino-acid flags with MHCflurry, with NetMHCpan EL, and with NetMHCpan BA.

At the primary cumulative ≤2% threshold per 100 candidate-specific opportunities, the mixed plan had lower paired MHCflurry burden than each of the five frozen comparators. NetMHCpan EL supported lower paired burden than AAY, GPGPG, no linker, and the selected uniform plan, but not GSG. NetMHCpan BA supported lower burden for the mixed plan than for no linker, AAY, and the selected uniform plan, while favoring GSG and GPGPG over the mixed plan. Supplementary Table S4 reports the estimates, nominal intervals, and Holm-adjusted tests for all 15 primary contrasts.

## Discussion

CAI-Vax addresses two construct-design decisions that remain after antigen selection: how to arrange antigens and intervening linkers, and how to encode the resulting protein cassette as a synonymous nucleotide sequence. HLA-aware search reduced MHCflurry-predicted junctional binding relative to amino-acid-pattern optimization.

Because MHCflurry also defined the search objective, this result shows that the search procedure successfully reduced its specified computational endpoint. It does not independently establish improved antigen processing, immunogenicity, or vaccine efficacy.

The comparison with exact and bounded solutions showed that local search decisions produce suboptimal complete constructs. Greedy search selects the next antigen using the immediate junction cost, but a locally favorable choice can restrict the quality of later junctions. The difference between single-start greedy search and the best result selected for each patient across all tested heuristics supports the use of exact optimization when computationally feasible and explicit lower bounds for larger instances. The combined result should not be interpreted as the performance of a single deployable algorithm. Adaptive search provides one practical alternative because it can stop using improvement history and a combinatorial lower bound without consulting the offline reference.

Interpretation of the junctional-binding endpoint also depends on its peptide definition. An eligible peptide was required to contain residues from both antigens flanking a junction. Changing linker length can therefore alter which peptides satisfy the endpoint, rather than merely changing their predicted binding ranks. A lower count should be interpreted as lower burden among the peptides defined by this rule, not as evidence that all possible junction-associated presentation has been reduced. Within-antigen peptides, linker-only peptides, and peptides containing residues from only one flanking antigen were not evaluated.

MHCflurry and NetMHCpan produced strong pooled correlations across the 186 frozen patient–plan observations, but agreement was weaker after accounting for patient-level differences and when selecting the best plan for an individual patient.^22,23^ This distinction matters because a predictor can rank patients similarly by overall burden while ranking alternative constructs differently within a patient. Rescoring fixed plans avoided improving a plan specifically for the second predictor, but agreement between two computational predictors is still a robustness assessment rather than biological validation.

The regret analysis quantified one consequence of this disagreement, although its scale requires care. Plans were selected using raw MHCflurry counts and evaluated using NetMHCpan EL counts per 100 candidate-specific opportunities. Regret may therefore reflect both predictor disagreement and differences in the opportunity denominator among plans. Reporting rankings under multiple predictors and scales is more informative than treating one model’s minimum as a predictor-independent optimum.

Synonymous optimization produced a similarly objective-dependent result. The repeat-aware encoder reduced maximum repeated 10-mer multiplicity in every patient while producing measurable losses in mean codon-use preference. Its performance did not generalize uniformly across repeat lengths: at six nucleotides, it performed worse than matched random encoding. In addition, the search-work threshold was calibrated using the historical 30-patient inputs and retained for the corrected rerun, even though objective weights were selected by leave-one-cohort-out comparison. These findings support describing the encoder as a retrospective internal analysis of sequence trade-offs.

More broadly, the results favor separating structural requirements from comparative objectives. Preservation of the intended protein and reading frame can be tested directly, whereas predicted HLA binding, codon use, repeat content, nucleotide composition, and RNA folding are model- or metric-dependent. Reporting these components separately keeps the trade-offs visible and avoids implying that a single composite score represents overall biological quality.

### Limitations

This study used fixed antigen sets from four retrospective published cohorts. It did not evaluate upstream antigen selection, and its findings do not establish superiority over external vaccine-design software. The HLA analysis was limited to predicted class I binding by peptides that contained residues from both antigens flanking a junction. It excluded within-antigen, linker-only, and single-antigen linker-edge peptides and did not evaluate antigen processing, natural presentation, T-cell recognition, class II responses, or clinical efficacy.

MHCflurry defined the primary optimization objective, so improved MHCflurry burden was not independent validation. NetMHCpan rescoring tested model dependence but did not provide an experimental reference. Historical records did not preserve the exact suffix build and model-file checksums of the original NetMHCpan installation. A separately documented 4.2e installation reproduced the retained counts, but it cannot establish the identity of the historical model files.

Cross-predictor regret combined raw-count selection with candidate-normalized evaluation. The repeat-aware encoder used a globally calibrated work threshold, so the leave-one-cohort-out weight selection did not constitute nested or independent validation. Sequence-feature flags, codon-use preferences, and repeat metrics are engineering proxies. No mRNA expression, stability, innate immune activation, protein production, synthesis yield, safety, or patient benefit was measured. Conclusions are therefore limited to the reported computational objectives, frozen model outputs, and internal baselines.

### Future studies

Experimental comparison of synonymous mRNAs encoding the same protein cassette could determine whether the reported repeat and codon-use trade-offs affect RNA stability, protein expression, innate immune sensing, or synthesis performance. Testing several synonymous sequences from the reported trade-off set would be more informative than comparing only one optimized sequence with one baseline.

For antigen ordering, immunopeptidomic analysis could test whether predicted junction-spanning peptides are naturally presented. T-cell assays could then assess whether presentation produces functional recognition. These experiments should include peptides excluded by the current both-antigen endpoint so that the effect of linker geometry can be evaluated directly.

Prospective evaluation should define the construct-selection rule before applying it to new cohorts. Such studies could compare single-predictor selection with consensus, ensemble, or robustness-based rules and evaluate whether cross-predictor stability predicts experimental outcomes. Direct comparison with existing construct-design software under shared antigen sets, endpoints, and computational budgets would also establish whether CAI-Vax offers advantages beyond its internal baselines.

## Conclusion

CAI-Vax is a computational workflow for arranging selected antigens, assigning linkers, and generating synonymous coding sequences. It reduced the junctional-binding objective used during search, quantified the limitations of heuristic construct ordering, and identified a trade-off between repeated-sequence reduction and codon-use preference. Rescoring fixed plans showed that the preferred construct could change with the HLA predictor, while synonymous-encoding results depended on the repeat length and sequence metric. These findings define the workflow’s computational behavior but do not establish biological or clinical superiority. Experimental and prospective studies are needed to determine whether the reported differences improve vaccine production or immune responses.

## Supporting information

Supplementary Methods and Results (including Tables S1-S8)

## Author contributions

Conceptualization, Methodology, Formal analysis, and Writing – original draft: Jalen Cai. Data curation and Investigation: Jalen Cai, Yulin Chen, Lareina Yuan, Eileen Liu, Yi Sun, and IpYan He. Software: Jalen Cai and Yulin Chen. Project administration: Jalen Cai, Wilson Ho-Man Lee, and Zhipeng Lu. Resources and Supervision: Wilson Ho-Man Lee and Zhipeng Lu. Validation: Jalen Cai, Wilson Ho-Man Lee, and Zhipeng Lu.

Visualization: Jalen Cai and Wilson Ho-Man Lee. Writing – review and editing: Wilson Ho-Man Lee and Zhipeng Lu.

## Acknowledgments

We thank the USC SC-ACE program, the CAI Foundation, and the USC Alfred E. Mann School of Pharmacy and Pharmaceutical Sciences for supporting this research experience.

## Use of AI-assisted tools

OpenAI ChatGPT/Codex and Anthropic Claude assisted with CAI-Vax software development. ChatGPT/Codex also assisted with manuscript editing, figures and statistical recalculation, as detailed in Supplementary Methods S1. The authors remain responsible for verifying the AI-assisted code, analyses, references and text and for the final content.

### Code and data availability

The versioned code and data archive for this study is https://doi.org/10.5281/zenodo.23489301 (CAI-Vax publication materials, version 1.0). It contains the analyzed source code based on commit b0585637a6b68b15bd22c84a5b37f26c56895ca7 (base software version 0.4.0.dev0) with the supplied patient-ID correction patch, processed GBM, HCC and PGV001 inputs, analysis tables, figures, data dictionary, reconstruction instructions and SHA-256 checksums. The base commit alone represents the earlier 30-patient inputs. The maintained development repository is https://github.com/JalenCai/CAI-Vax; the archived source provides reproducibility without requiring access to that repository.

Exact processed GBM, HCC and corrected PGV001 inputs are included under their source-specific terms with citations and modification notes. The processed melanoma table is not redistributed; the archive provides the exact publisher-source URLs, harmonization procedure and expected processed-file hash so readers can reconstruct it locally. All eleven source supplements were anonymously downloaded and matched by SHA-256, and the supplied reconstruction helper reproduced all four exact processed input hashes. Licensed predictor installations, model assets and raw sequence-bearing prediction caches are excluded.

The historical CAI-Vax_Separate_Reproduction_Full_20261010.zip documents new predictor installations and sequence-free comparisons for the earlier 180 frozen plans. The current correction records separately document the two newly scored patients and 186-row merged analysis. NetMHCpan must be obtained from DTU under its license. Supplementary Methods distinguish the historical reproduction from the current corrected analysis.

