## Supplementary Methods and Results (including Tables S1-S8) for "CAI-Vax: Computational Design of Personalized mRNA Cancer Vaccine Constructs After Antigen Selection"

CAI-Vax Supplementary Methods and Results

Publication package version 1.0 19 October 2026

This supplemental document defines the computational endpoints, search methods, synonymous-encoding procedures, statistical calculations, and retained results. The accompanying package supplies the source tables, source archive, separate predictor reproduction, data dictionary, and provenance records. Conclusions concern the stated model and sequence metrics.

### S1. Analysis materials and reproduction boundaries

The historical analysis source is commit b0585637a6b68b15bd22c84a5b37f26c56895ca7, package version 0.4.0.dev0. The accompanying source archive preserves this snapshot. Detailed provenance records are supplied separately in the archive.

The versioned code and data archive for this study is https://doi.org/10.5281/zenodo.23489301 (CAI-Vax publication materials, version 1.0). It contains the analyzed source code based on commit b0585637a6b68b15bd22c84a5b37f26c56895ca7 (base software version 0.4.0.dev0) with the supplied patient-ID correction patch, processed GBM, HCC and PGV001 inputs, analysis tables, figures, data dictionary, reconstruction instructions and SHA-256 checksums. The base commit alone represents the earlier 30-patient inputs. The maintained development repository is https://github.com/JalenCai/CAI-Vax; the archived source provides reproducibility without requiring access to that repository. Melanoma is supplied through the verified publisher-source retrieval and reconstruction route. Third-party input terms and attribution are documented in the archive; licensed predictor installations, model assets and raw sequence-bearing predictions are excluded.

CAI-Vax_Separate_Reproduction_Full_20261010.zip is included under reproducibility. It records configurations, execution commands, predictor and model checksums, dependency provenance, and sequence-free count and statistical comparisons. The source-file catalog and figure/table source map identify the numerical inputs to each display.

The accompanying source tables report 128 HLA and 150 frozen-predictor paired contrasts recalculated with exact conditional signed-rank tests, together with four unadjusted cohort repeat tests. Integer-count rational arithmetic determines differences and ties. Historical statistical outputs and the earlier source/reproduction archives remain separately identified. S11 gives the review calculation and nominal interval procedures.

OpenAI ChatGPT/Codex and Anthropic Claude were used to assist with development of the CAI-Vax software. ChatGPT/Codex also assisted with manuscript and supplement comparison and editing, figure revisions, and preparation and checking of the statistical recalculation. The authors remain responsible for verifying the AI-assisted code, analyses, references and text and for the final content.

### S2. Cohort inputs and harmonization

Each patient is identified by (dataset_id, patient_id). The processed files contribute GBM 4/36, HCC 9/107, melanoma 6/98 and PGV001 12/116 evaluable patients/antigens, totaling 31/357. GBM was the development cohort; the other three were locked before historical retrospective evaluation. Review found that Table S3 labels one PGV001 block PGV-019, whereas Table S2 uses PID-019. The corrected adapter normalizes both prefixes to PID before the HLA join. The former parser had carried PID-017 across ten PID-019 peptide rows. The correction yields separate ten-peptide patients with their distinct source HLA metadata. Of 13 source IDs, PID-002 remains excluded because its entries lack extractable peptide sequences. HCC1395 remains an excluded cell-line example.

Each processed table retains the cohort and patient identifiers, antigen identifiers, source metadata, and the antigen sequence field (long_peptide) used by the cohort loader. Sequences are stripped, uppercased, and required to be nonempty strings of standard amino-acid symbols. The input-access inventory records exact file hashes and included counts. Adapters retain upstream order and available selection/HLA metadata; the archived source defines their broader schemas. Invalid or missing sequences and mapping failures are reported. No patient’s antigens are pooled with another’s. Duplicate-ID handling is recorded; exact processed files and hashes define this study.

### S3. Amino acid flags and intentional linker repeats

The component feature window is the last five residues of the left antigen, the linker, and the first five residues of the right antigen; short antigen flanks use available residues. A component flag is the logical union of the following four tests. Count represents the number of flagged junctions.

| **Feature** | **Exact rule** |
| --- | --- |
| Identical residue run | Maximum consecutive identical-residue run ≥5 |
| Basic residue run | Consecutive residues from K, R, H ≥5 |
| Hydrophobic residue run | Consecutive residues from A, I, L, M, F, V, W, Y ≥5 |
| Low complexity | Window length ≥5 and number of distinct residues ≤2 |
| Repeat annotation | A junction-crossing amino-acid 4-mer occurs more than once by Python cassette.count; this counts nonoverlapping matches |
| Intentional shared linker repeat | The same linker name is reused at another seam; shared-linker repeats are annotated separately and excluded from the nonshared-repeat objective |

The shared-repeat classification uses linker-name reuse. This annotation rule can therefore be broader than a sequence-specific attribution of repeat origin.

The corrected amino-acid comparator minimizes additive flagged-junction count. Per-edge ties use shorter linker length, then no linker, GSG, AAY, GPGPG. Source-order uniform GSG, no-linker and AAY baselines in the threshold grid differ from optimized-order uniform plans in the frozen six-plan rescoring.

Corrected scalar search used exact search for n≤14. Larger cases compared matched configurations: 32 multistart 2-opt restarts and beam widths 16, 32, 64, 128 for both amino-acid and HLA objectives.

### S4. Peptide endpoint and normalization

Lengths are 8, 9, 10, and 11 amino acids. Each included peptide must contain at least one residue from both antigens flanking its junction and spans the complete intervening linker if present. Single-antigen linker-edge peptides, linker-only peptides, and within-antigen peptides are outside this endpoint. HLA alleles are source-provided class I alleles checked for predictor support; no allele imputation or class II endpoint was used.

Within each junction, duplicate peptide strings are collapsed before multiplying by supported alleles. MHCflurry inference can cache a construct-wide union of peptide–allele pairs, but the burden numerator sums qualifying pair occurrences at each junction. Thus, the same qualifying peptide–allele pair at two junctions can contribute twice. NetMHCpan frozen rescoring instead sums qualifying pairs in the construct-wide union. The denominator for per-100 reporting is the construct-wide unique eligible peptide count multiplied by the supported allele count. Consequently, the MHCflurry per-100 value is an occurrence-to-unique-opportunity ratio.

The current analysis contains 186 frozen plans: 174 unchanged plans with historical fingerprints and 12 newly reconstructed plans for corrected PID-017 and PID-019. Two plans repeated eligible peptides across junctions: melanoma patient 1 uniform AAY had 264 within-junction peptide occurrences versus 260 unique peptides; patient 4 uniform GPGPG had 182 versus 181. Their supported allele counts were three and one. Both had zero recorded MHCflurry counts at ≤0.5% and ≤2%; the other 184 plans had no cross-junction duplicates. Since contributions are nonnegative, these zeros establish that union deduplication changes none of the frozen counts or derived comparisons at these two thresholds. This is a statement about recorded outputs and these constructs. The duplicate-proof tables preserve the counts and source rows.

Threshold counts use numeric percentile rank ≤t, for t=0.5, 1, 2, 5. The ≤2% count includes the ≤0.5% strong-binder count. Raw is the summed occurrence count; per junction divides by the number of antigen seams; per allele divides by supported class I allele count; per 100 opportunities multiplies raw by 100 and divides by the plan-specific union peptide–allele denominator. Exact boundary values are included. DTU displayed SB uses rank <0.5 and WB uses rank <2 excluding SB.

Longer linkers can remove peptides from this both-antigen endpoint by changing window geometry. NetMHCpan threshold categories and numeric-rank definitions are documented at https://services.healthtech.dtu.dk/services/NetMHCpan-4.2/.

### S5. Predictors and frozen plan definitions

MHCflurry records specify version 2.2.1 and affinity percentile ranks.^22^ NetMHCpan EL and BA ranks were parsed separately.^23^ Input was a peptide list; the wrapper used -BA -p -inptype 1 with the patient allele and peptide file.

The historical reproduction used documented new MHCflurry 2.2.1 and NetMHCpan 4.2e installations. Current correction inference uses the same public MHCflurry model hashes and the local licensed NetMHCpan 4.2e installation. The current Torch runtime is 2.14.1; all runtime versions, commands and hashes are recorded. For 4,800 common peptide-allele pairs, old and new MHCflurry predictions give identical classifications at all four thresholds.

#### *Separate predictor reproduction on 10 October 2026*

The 10 October reproduction reconstructed 180 frozen plans for the historical 30-patient inputs and matched all 720 MHCflurry and 1,440 NetMHCpan EL/BA threshold counts. Thirty mixed-plan fingerprints matched original references; the other 150 references came from the reconstruction audit. A corrected amino-acid comparator was replayed.

The separate installations and count comparisons are summarized below. Model, installer, configured-wrapper, executable, and model/data hashes and complete dependency records are supplied in the reproduction archive. No context, pathogen or neoepitope mode was enabled.

| **Predictor** | **New installation and mode** | **Retained count comparison** |
| --- | --- | --- |
| MHCflurry 2.2.1 | Public ten-network pan-allele affinity ensemble models_class1_pan selected.20200610; affinity percentile ranks from peptide lists. | 720/720 matched for historical inputs; current correction separately records 12 new plans |
| NetMHCpan 4.2e | Peptide-only input with -BA -p -inptype 1; EL and BA ranks parsed separately. | 1,440/1,440 EL/BA matched for historical inputs; current correction separately records 12 new plans |

The current corrected analysis contains 186 plans: 174 unchanged plans from 29 patients and 12 newly reconstructed and scored plans for PID-017 and PID-019, with complete supported-allele predictions. All affected searches and all 31 repeat-aware encodings were rerun. All patient summaries and 282 signed-rank tests were recomputed. Model hashes, commands and count provenance are supplied in the archive; detailed correction history is in the review record.

Patient-cluster correlations use 2,000 resamples, seed 472026; primary EL regret uses 2,000, seed 522026. The historical reproduction and current correction are separate evidence records. No comparison to historical outputs establishes the identity of original historical predictor installations.

| **Frozen plan name** | **Operational definition** |
| --- | --- |
| Mixed-linker | Recorded HLA-minimizing mixed-linker plan with its optimized antigen order and seam linkers |
| Selected uniform | Recorded minimum among the four uniform-linker optimized-order plans |
| No linker | Recorded optimized antigen order with no linker at all seams |
| Uniform GSG | Recorded optimized antigen order with GSG at all seams |
| Uniform AAY | Recorded optimized antigen order with AAY at all seams |
| Uniform GPGPG | Recorded optimized antigen order with GPGPG at all seams |

There are six rows per patient even when distinct plan labels yield identical sequences or scores. Minima and tie-aware plan sets use all six labels. Construct fingerprints define identity of the assembled protein. Complete frozen tables report 186/186 supported-allele rows.

### S6. Scalar search and certificates

For antigens i and j, c(i,j)=min over linkers of the chosen scalar edge cost. Held–Karp state D(S,j) stores the minimum cost of a path visiting exactly S and ending at j. Initialize D({j},j)=0. Extend by D(S∪{j},j)=min over i∈S of D(S,i)+c(i,j), retaining a path predecessor; the answer is min over j of D(V,j). This is a Hamiltonian path and is exact for the declared edge-additive objective up to n=14.^24^

The larger-instance custom MILP adds a zero-cost depot, binary directed arcs, one incoming and outgoing arc per node, and Miller–Tucker–Zemlin ordering constraints to eliminate antigen-only subtours.^25^ SciPy milp invokes HiGHS.^26^ The corrected certification/recovery time limit is 30 seconds and mip_rel_gap=0. The adaptive offline reference uses a separate 10-second MILP limit. A valid incumbent without a certificate is best feasible; a combinatorial lower bound remains a bound.

Single-start greedy starts at the lexicographically smallest antigen ID; all-start greedy starts at every antigen and keeps the best. Multistart 2-opt uses 32 random restarts, seed 12345, and strictly improving path reversals. Beam widths are 16, 32, 64, and 128. On 25 exact instances, success counts were single-start 5, all-start 7, multistart 2-opt 16, and beam 11/14/14/15 by width. The per-patient union reached 19/25 and missed six. This union is an offline best across methods. Five of six larger instances were certified. Melanoma patient 2 (17 antigens) had a best valid MILP incumbent of 4, heuristic score 6, combinatorial lower bound 0, and uncertified additive gap 4.

The predecessor comparator was exhaustive order search for n≤7 and lexicographically-smallest-start greedy for larger cases, both with per-edge optimal linker choices. It is identified by this configuration. Comparing its frozen scores with corrected search gives 21/31 improvements, mean reduction 4.52, maximum 17. Model-inference time and path-search time are separate costs.

### S7. Pareto recurrence and label limits

The custom multiobjective state is the visited-antigen subset and last antigen. Each label stores cumulative amino-acid flags and HLA binder occurrences, plus the full order/linker path. Singleton labels start at zero. Each unused antigen and each linker extends the path by the two edge costs. Dominated labels are removed; equal vectors retain one representative path. Linkers are carried in the path payload.

Uncapped label setting is exact within n ≤ 11. Capped search uses NSGA-II crowding distance, preserving objective extremes when possible and deterministic tie ordering.^27^ Real-patient coverage used caps 8, 16, 32, 64. Synthetic n=9 cap tests used 4, 6, 8, 16, 32 and seeds 0–3; exhaustive validation used n=4–6 and seeds 0–2 plus feasible real cases n ≤ 5. Objective-vector coverage, additive epsilon, and hypervolume quantify approximation loss.

### S8. Adaptive stopping and measured work

The deployed algorithm first computes a combinatorial lower bound and a greedy incumbent. Random restart batches are (1,1,2,4,8,16), followed by counted strictly improving 2-opt. It stops after a batch if best score ≤ lower bound, gap ≤ 0, two consecutive batches have no strict incumbent improvement, or elapsed wall time exceeds 5 seconds. Base seed is 12345; eight-seed diagnostics use 12345+1009s for s=0…7. The exact reference is called only afterward.

The current table combines 29 retained base-seed adaptive runs with newly computed runs for corrected PID-017 and PID-019. It records starts, counted order evaluations and path-search time separately from predictor inference. The schedule permits at most 32 random restarts plus one greedy start.

Adaptive search ran for all 31 patients. Certified-reference evaluation used 30, omitting only melanoma patient 2 because the offline reference was uncertified. Of 16 runs matching a certified reference, 11 stopped by attaining their deployed lower bound; the 11 are a subset of the 16. The omitted case's reference is labeled as best known.

### S9. Synonymous encodings random draws and quality control

The intended protein is the assembled cassette, prepended with M only if it does not already begin with M. Encoders preserve that intended protein and add the fixed terminal stop-codon convention TAA/TGA used by the implementation. Standard encoders include the human-codon-use CAI-Vax heuristic, highest-frequency codon, GC-balanced, low-repeat, and uniform random synonymous codons. Its per-codon lookup values and terminal stops are included by the metric implementation.

The standard balanced encoder greedily minimizes, for each appended synonymous codon, 1.0×absolute cumulative GC-fraction deviation from 0.52 +1.0×new CpG count +0.5×new TA count +1.0×max(0, trailing homopolymer run−3) −0.3×codon-use preference. Highest-frequency minimizes negative codon preference; GC-balanced minimizes cumulative GC deviation; the baseline named low_repeat actually minimizes the appended homopolymer run. Low-CpG and low-UpA generator variants minimize newly created CG and TA respectively. Greedy ties are ordered by higher codon preference then codon string; a near-tie check uses 10⁻⁹. Random encoding alone samples uniformly from synonymous codons. Default stop strategy tandem_stop_2 uses primary TAA followed by TGA; translation halts at the first stop. Full codon lookup tables remain in the archived source. The original composite benchmark selected among 12 internally generated CAI-Vax designs using its configured composite score; this selection is part of the retrospective comparison.

The composite benchmark generated 500 uniform synonymous random encodings per patient using base seed 12345 and random.Random(seed+i). The repeat-length comparison used 120 matched-protein encodings. Initial methionine and terminal stops followed the same conventions. Optional folding used ViennaRNA when available; unavailable results remained missing.^28^

Higher codon-use preference and higher composite score are favorable; lower maximum repeated-k-mer multiplicity, repeat burden, and homopolymer length are favorable. GC percentage is descriptive without a specified target; absolute GC deviation is minimized only in a configured target objective. CpG/UpA density and folding energy do not have a universal biological favorable direction. Their sign in a configured objective is given explicitly below. Composite exceedance P=(1+number of random scores ≥ candidate score)/(N+1) is one-sided. Its empirical score percentile uses strict <. Component percentiles use raw metric ≤. Wins against the random median require strict favorable improvement; ties are excluded from the one-sided patient sign-test denominator. Two-sided patient Wilcoxon tests remain separate.

Structural QC checks DNA alphabet A/C/G/T, coding length divisible by three, expected translation, allowed terminal stops and internal stops, and a configured maximum sequence length of 12,000 nt. Default feasibility warnings include global GC outside 30–75%, 50-nt window GC outside 20–85%, homopolymer length above 10, and nucleotide 10-mer multiplicity above 3. Missing optional models are recorded as unavailable. The repeat rerun used validator version 1.0.0; all 31 outputs passed structural validation and protein preservation.

Initiating methionine was added to 29 of 31 source-order GSG benchmark cassettes; HCC patients N09 and N31 already began with M. The sequence-free methionine audit records per-patient flags and source/intended-protein hashes.

### S10. Repeat objective configuration and held out evaluation

The comparator is the starting standard CAI-Vax source-order GSG coding sequence, encoded with seed 12345. The repeat encoder preserves its intended protein. Maximum nucleotide k-mer multiplicity counts overlapping windows. Primary k=10; sensitivity lengths are 6,8,10,12,15,20. Lower maximum multiplicity is favorable.

Minimized objective: 1×maximum 10-mer multiplicity + 0.25×sum of repeated-10-mer excess counts + 0.20×max(0, maximum homopolymer−6) + 0.02×absolute GC percentage deviation from 52 + 0.05×100×CpG fraction + 0.02×100×UpA (DNA TA) fraction − 0.50×mean human codon-use preference. The synonymous hill climb tests alternatives in deterministic position/codon order, uses strictly improving objective moves with tolerance 10^-9^, and breaks candidate ties by objective, position, and alternative codon. Hard constraints are GC within 30–75% inclusive, maximum homopolymer ≤ 10 and no greater than the starting value, codon-use preference ≥ starting preference−0.02, and identical intended translation. Maximum iterations is 400. The work threshold is 4,000 objective evaluations, checked after a complete sweep, so it can be exceeded. Attempted substitutions and objective evaluations are separate counters, with codon-prefiltered alternatives not all reaching objective evaluation.

Candidate configurations are frozen (repeat weight 1), w_repeat_2x (weight 2), and w_repeat_4x (weight 4); other weights are unchanged. For each held-out cohort, eligible configurations have training-cohort mean codon-use loss <0.02. Selection prefers eligibility, then largest mean k=10 reduction rounded to nine decimal places, then the explicit priority frozen, 2x, 4x. If none were eligible the code would still choose its best-ranked candidate. All four folds selected frozen. Each patient belongs to the held-out set in exactly one cohort fold; every patient refers to all 31 held-out patients aggregated across folds.

The historically calibrated candidate work thresholds were 2,000, 4,000, and 8,000. The rule selected the smallest zero-failure threshold within 0.2 of the largest observed mean repeat reduction. The largest mean was 5.9667 and the selected 4,000 threshold yielded 5.7667. This work calibration used all cohorts in the former 30-patient dataset. The selected 4,000 threshold was held fixed in the corrected 31-patient rerun; only objective-weight selection was isolated by cohort. The result is retrospective internal evaluation.

A selected-configuration rerun in all 31 patients recomputed repeat reduction and codon-use loss; the 29 unaffected cases exactly matched earlier sequence hashes and deterministic work counts. Actual objective counts were 4,004–5,277 and accepted moves 13–29. The per-patient work table supplies hashes, attempted substitutions, validation, stop reasons, and repeat-length results.

### S11. Statistical estimators intervals and families

For paired differences, the Hodges–Lehmann estimate is the median of the Walsh averages, including self-pairs.^29^ Estimates and interval endpoints in the review recalculation use the rational patient differences described below. For n≥6, the procedure sorts N=n(n+1)/2 averages and uses signed-rank mean μ=n(n+1)/4 and variance σ²=n(n+1)(2n+1)/24. The lower zero-based index is k=floor(μ−zσ), where z is the 97.5th percentile of the standard normal distribution, clipped into the array. Bounds are Walsh[k] and Walsh[N−1−k]. For n=31, there are 496 averages and zero-based indices 147 and 348. This retained nominal interval uses a normal approximation without tie-corrected or exact inversion. For n<6, it is the minimum-to-maximum Walsh range.

Two-sided Wilcoxon signed-rank P values were recalculated after review from the exact conditional distribution of the positive rank sum.^30^ Patient-level normalized values were formed as exact rational numbers from integer numerators and recorded denominators, and differences were subtracted before ranking. True zero differences were discarded; equal absolute nonzero differences received average ranks. Ranks were doubled to integers, and a dynamic program counted all positive/negative sign assignments. The two-sided probability was twice the smaller inclusive tail probability, capped at 1; all-zero differences received P=1. No Monte Carlo approximation was used. Validity requires independent patient differences and sign symmetry under the null. This update retained the 128 HLA and 150 frozen-predictor contrast definitions and their original Holm families, plus the four unadjusted repeat-cohort tests. Rank-biserial correlations exclude zeros.

| **Quantity** | **Resampling or test specification** |
| --- | --- |
| Primary HLA HL interval | Nominal Walsh normal-approximation interval; no bootstrap |
| Frozen normalized paired contrasts | Nominal Walsh intervals; exact conditional signed-rank P values with tied ranks and zeros handled explicitly |
| Pooled predictor correlations | Patient-cluster percentile bootstrap, 2,000 resamples, seed 472026 |
| Pessimistic EL regret median and HL | Patient percentile bootstrap, 2,000 resamples, seed 522026 |
| Mean codon-use loss | Patient-mean percentile bootstrap within cohort, 2,000 resamples, seed 12345 |
| Regret driver numeric tests | Spearman permutation tests, 20,000 permutations, seed 532026 |
| Patient proportions | Clopper–Pearson two-sided 95% intervals |

Bootstrap sampling draws patients with replacement and retains their correlated plans together.^32^ For frozen regret each patient first contributes one scalar regret. The primary HLA Holm family contains eight named contrasts at one threshold and normalization (16 separate eight-test families, 128 rows). It is not one family covering the 16 c1 sensitivity cells. Frozen predictor paired comparisons form five-test families at each predictor endpoint and normalization (30 families, 150 rows). The primary weak/2%/per-100 table comprises three five-test families. The nine exploratory regret drivers form one Holm family. The four frozen cohort repeat tests are unadjusted. The codon-loss table reports bootstrap mean intervals and margin checks.

The eight HLA contrasts are corrected amino-acid mixed minus HLA mixed; amino-acid uniform minus HLA uniform; source-order GSG, AAY, and no linker each minus HLA mixed; selected HLA uniform minus HLA mixed; predecessor mixed minus corrected mixed; and predecessor uniform minus corrected uniform. A positive effect favors the second plan. Machine IDs and family membership are retained in the data dictionary.

### S12. Within patient ranking and cross predictor regret

Spearman correlation is Pearson correlation of average tied ranks.^36^ Within-patient correlations require at least three finite rows and nonconstant scores in both predictors. Thirty patients have defined MHCflurry–EL and MHCflurry–BA correlations: EL median 0.3459 (IQR −0.0735–0.5658) and BA median −0.1328 (IQR −0.3529–0.1547). Missing correlations are excluded from these summaries. Patient-centered pooling subtracts each patient’s arithmetic mean from each predictor’s six scores, then computes Spearman across 186 residual pairs; constant patient residuals can remain. This gives EL 0.2824 and BA −0.1910, compared with raw pooled values 0.8714 and 0.7190. No explicit within/between-patient variance decomposition or causal contribution calculation was done.

MHCflurry selection uses raw cumulative ≤2% counts, with minima across all six candidates and exact-equality ties (tolerance zero). For each candidate, EL evaluation is its own cumulative count divided by its own eligible unique peptide-allele opportunity denominator and multiplied by 100. Pessimistic regret=max EL over the MHCflurry minimum set−min EL over all six plans. These different selection/evaluation scales are intentional. The best-set function drops missing scores and returns an empty set only when no finite values remain. Every retained patient had six finite candidate scores, so observed minima used all six. Zero regret occurs in 10/31; its proportion is 0.3226 with Clopper–Pearson interval 0.1668–0.5137. Median regret=0.210831; HL=0.237323; maximum=1.422469 (PGV001 PID-012). The HL patient-bootstrap interval is 0.126262–0.401070.

Nine exploratory drivers assessed antigen count, junction count, supported HLA count, focus-plan opportunities, cohort, tie frequency, mean best-to-second margin, search certification, and construct length. Numeric/binary variables used two-sided permutation Spearman tests (20,000 permutations, seed 532026, plus-one correction) and Theil–Sen slopes; slope intervals used 2,000 paired-patient bootstrap resamples, seed 522026.^34^ Cohort used Kruskal–Wallis.^35^ The archived dictionary defines the recorded margin and certification fields. None survived the nine-test Holm family (smallest raw P about 0.00620; adjusted about 0.05580).

Best-set equality is exact equality of tied sets of six plan labels (6/31 EL); overlap means nonempty set intersection (19/31). The 14/31 common-non-dominated result asks whether at least one label is non-dominated in each of three separate views: (AA flags,MHCflurry), (AA flags,EL), and (AA flags,BA).

### Supplementary Results

All effects concern fixed patient candidates and numerical outputs. The primary amino-acid-versus-HLA contrast favored HLA search in all 16 sensitivity settings (Table S1). All 31 patient differences were positive at the primary ≤2% per-junction setting; some sensitivity settings included ties or negative differences. Other contrasts include null and threshold-dependent reversals. The full 128 HLA and 150 predictor contrasts accompany this supplement.

### Table S1. Primary contrast across the complete sensitivity grid

| **Rank** | **Scale** | **HL** | **Nominal 95% interval** | **Holm P** | **Positive / zero / negative** |
| --- | --- | --- | --- | --- | --- |
| 0.5% | Raw | 7.5000 | 5.0000–10.0000 | 1.043e-07 | 27/4/0 |
| 1% | Raw | 16.2500 | 11.5000–21.0000 | 7.451e-09 | 31/0/0 |
| 2% | Raw | 33.5000 | 25.0000–41.5000 | 7.451e-09 | 31/0/0 |
| 5% | Raw | 45.7500 | 28.5000–61.5000 | 1.304e-08 | 30/1/0 |
| 0.5% | Per junction | 0.8056 | 0.5556–1.0497 | 1.043e-07 | 27/4/0 |
| 1% | Per junction | 1.7500 | 1.2051–2.2500 | 7.451e-09 | 31/0/0 |
| 2% | Per junction | 3.4919 | 2.7479–4.4798 | 7.451e-09 | 31/0/0 |
| 5% | Per junction | 4.7212 | 2.9000–6.6368 | 1.304e-08 | 30/1/0 |
| 0.5% | Per allele | 2.2500 | 1.5000–3.0833 | 1.043e-07 | 27/4/0 |
| 1% | Per allele | 4.6667 | 3.5833–5.8333 | 7.451e-09 | 31/0/0 |
| 2% | Per allele | 9.6667 | 7.8333–12.4167 | 7.451e-09 | 31/0/0 |
| 5% | Per allele | 13.5000 | 10.0000–16.9167 | 1.304e-08 | 30/1/0 |
| 0.5% | Per 100 opportunities | 0.6199 | 0.4386–0.7898 | 2.086e-07 | 26/4/1 |
| 1% | Per 100 opportunities | 1.2850 | 1.0314–1.5467 | 7.451e-09 | 31/0/0 |
| 2% | Per 100 opportunities | 2.7929 | 2.3605–3.1782 | 7.451e-09 | 31/0/0 |
| 5% | Per 100 opportunities | 2.2568 | 1.4596–2.9770 | 5.051e-06 | 26/0/5 |

Contrast is corrected amino-acid-pattern minus HLA-aware mixed-plan burden. Positive favors HLA-aware. The all-31 statement applies to the primary ≤2% per-junction setting; direction counts are supplied separately for every cell. All 128 threshold-grid contrasts with raw and adjusted tests are in supplement_hla_all_128_contrasts.tsv.

All 31 synonymous outputs passed the recorded structural checks. Table S2 gives the complete per-patient work accounting. The 4,000-evaluation threshold was checked after a sweep; observed counts exceeded that threshold. Tables 2 and S3 retain the codon-use trade-off and the four-patient GBM uncertainty.

### Table S2. Repeated encoder observed evaluation counts

| **Cohort** | **Patient** | **Evaluations** | **Moves** | **k10 reduction** | **Codon-use loss** |
| --- | --- | --- | --- | --- | --- |
| GBM | 1 | 4,029 | 23 | 7 | 0.0073 |
| GBM | 2 | 4,460 | 13 | 2 | 0.0045 |
| GBM | 3 | 4,010 | 13 | 4 | 0.0054 |
| GBM | 4 | 4,532 | 13 | 3 | 0.0048 |
| HCC | N06 | 4,428 | 13 | 3 | 0.0035 |
| HCC | N09 | 4,203 | 21 | 8 | 0.0048 |
| HCC | N13 | 4,634 | 16 | 4 | 0.0045 |
| HCC | N22 | 4,062 | 18 | 4 | 0.0044 |
| HCC | N24 | 4,042 | 22 | 5 | 0.0048 |
| HCC | N25 | 4,780 | 19 | 7 | 0.0052 |
| HCC | N27 | 4,120 | 14 | 2 | 0.0065 |
| HCC | N30 | 4,140 | 16 | 6 | 0.0056 |
| HCC | N31 | 5,277 | 29 | 9 | 0.0049 |
| melanoma | 1 | 4,528 | 23 | 8 | 0.0071 |
| melanoma | 2 | 4,110 | 16 | 5 | 0.0032 |
| melanoma | 3 | 4,763 | 17 | 6 | 0.0035 |
| melanoma | 4 | 4,658 | 19 | 9 | 0.0040 |
| melanoma | 5 | 4,269 | 20 | 8 | 0.0038 |
| melanoma | 6 | 4,278 | 22 | 14 | 0.0041 |
| PGV001 | PID-006 | 4,051 | 16 | 6 | 0.0040 |
| PGV001 | PID-008 | 4,556 | 14 | 3 | 0.0038 |
| PGV001 | PID-010 | 4,361 | 15 | 5 | 0.0051 |
| PGV001 | PID-012 | 4,255 | 15 | 4 | 0.0041 |
| PGV001 | PID-014 | 4,157 | 16 | 5 | 0.0049 |
| PGV001 | PID-015 | 4,004 | 15 | 6 | 0.0037 |
| PGV001 | PID-016 | 4,548 | 15 | 5 | 0.0038 |
| PGV001 | PID-017 | 4,453 | 13 | 4 | 0.0034 |
| PGV001 | PID-019 | 4,693 | 16 | 5 | 0.0043 |
| PGV001 | PID-020 | 4,735 | 15 | 5 | 0.0037 |
| PGV001 | PID-022 | 4,506 | 15 | 6 | 0.0042 |
| PGV001 | PID-023 | 4,620 | 14 | 5 | 0.0046 |

New review-rerun counts for the selected frozen configuration. All 31 outputs passed current structural QC and preserved the intended protein. See the TSV for elapsed time, attempted substitutions, validator version, and sequence hashes.

### Table S3. Repeat effects and codon use trade offs

| **Cohort** | **n** | **Repeat HL** | **Interval type, bounds** | **Raw P** | **Mean codon loss** | **Bootstrap 95% interval** |
| --- | --- | --- | --- | --- | --- | --- |
| GBM | 4 | 3.75 | Walsh range 2–7 | 0.125 | 0.0055 | 0.0046–0.0067 |
| HCC | 9 | 5.5 | Nominal CI 3.5–7.5 | 0.003906 | 0.0049 | 0.0044–0.0054 |
| Melanoma | 6 | 8.0 | Nominal CI 5.5–11.5 | 0.03125 | 0.0043 | 0.0035–0.0055 |
| PGV001 | 12 | 5.0 | Nominal CI 4.5–5.5 | 0.0004883 | 0.0041 | 0.0039–0.0044 |

Review-recalculated estimates use the corrected 31-patient inputs; comparator is the starting standard CAI-Vax sequence. Codon losses use arithmetic human codon-use preference. GBM n=4 and raw P=0.125 are preserved. The codon-loss bootstrap was recomputed for the corrected cohort. All losses are measurable even while upper bounds satisfy the configured 0.02 margin; no equivalence conclusion follows.

Predictor transfer was incomplete. At cumulative ≤2% per 100 opportunities, EL supported the mixed plan against AAY, GPGPG, no linker and selected uniform, with no significant advantage against GSG. BA favored GSG and GPGPG over mixed. Across the HLA threshold grid, the primary mixed-versus-amino-acid contrast remained favorable and significant in all 16 settings, whereas mixed-versus-selected-uniform and AAY comparisons varied by threshold and scale. Repeat-aware encoding reduced maximum 10-mer multiplicity in all 31 patients but incurred measurable codon-use losses and performed worse than matched random encoding at k=6. GBM remained nonsignificant (n=4; P=0.125). The complete contrast tables preserve the null results and reversals.

### Table S4. Primary frozen predictor paired contrasts

| **Endpoint** | **Comparator** | **HL difference** | **Nominal 95% interval** | **Holm P** |
| --- | --- | --- | --- | --- |
| MHCflurry | No linker | -0.7318 | -0.9702 to -0.5608 | 1.863e-08 |
| MHCflurry | GSG | -0.5917 | -0.8247 to -0.4061 | 7.153e-07 |
| MHCflurry | AAY | -0.2302 | -0.3589 to -0.1515 | 1.907e-06 |
| MHCflurry | GPGPG | -3.2394 | -4.6638 to -2.2985 | 2.98e-08 |
| MHCflurry | Selected uniform | -0.1966 | -0.3365 to -0.1058 | 3.815e-06 |
| NetMHCpan EL | No linker | -0.5917 | -0.8071 to -0.3665 | 3.725e-08 |
| NetMHCpan EL | GSG | 0.0439 | -0.0823 to 0.1389 | 0.6102 |
| NetMHCpan EL | AAY | -0.5129 | -0.6759 to -0.3642 | 1.043e-07 |
| NetMHCpan EL | GPGPG | -0.4170 | -0.7083 to -0.2147 | 8.082e-05 |
| NetMHCpan EL | Selected uniform | -0.2872 | -0.4567 to -0.0973 | 0.0013 |
| NetMHCpan BA | No linker | -0.3172 | -0.6321 to -0.0672 | 0.01383 |
| NetMHCpan BA | GSG | 0.3568 | 0.1389 to 0.6784 | 0.003583 |
| NetMHCpan BA | AAY | -0.9028 | -1.1912 to -0.6170 | 1.304e-07 |
| NetMHCpan BA | GPGPG | 0.3968 | 0.1337 to 0.7193 | 0.006838 |
| NetMHCpan BA | Selected uniform | -0.5244 | -0.8523 to -0.2105 | 0.002982 |

Mixed-linker plan minus comparator; negative favors mixed. All are cumulative ≤2%, per 100 candidate-specific opportunities, n=31. Five comparisons form one Holm family per predictor. EL GPGPG favors mixed (HL -0.4170; nominal CI -0.7083 to -0.2147; adjusted P=8.082e-05), whereas EL GSG is not supported. BA GSG and GPGPG favor the uniform comparator. The zero-count proof in S4 establishes that construct-union deduplication leaves frozen MHCflurry counts unchanged at the two recorded thresholds. The full 150-contrast table supplies all endpoints and scales.
